# Does the Antisecretory Peptide AF-16 modulate fluid balance and inflammation in experimental peritonitis induced sepsis?

**DOI:** 10.1101/2020.04.14.040857

**Authors:** Annelie Barrueta Tenhunen, Jaap van der Heijden, Ivan Blokhin, Fabrizia Massaro, Hans Arne Hansson, Ricardo Feinstein, Anders Larsson, Jyrki Tenhunen

## Abstract

Sepsis is a life-threatening condition due to a dysregulated immunological response to infection. Apart from source control and broad-spectrum antibiotics, management is based on fluid resuscitation and vasoactive drugs. Fluid resuscitation implicates the risk of volume overload, which in turn is associated with longer stay in intensive care, prolonged use of mechanical ventilation and increased mortality.

Antisecretory factor (AF), an endogenous protein, is detectable in most tissues and in plasma. The biologically active site of the protein is located in an 8-peptide sequence, contained in a synthetic 16-peptide fragment, named AF-16. The protein as well as the peptide AF-16 has multiple modulatory effects on abnormal fluid transport and edema formation/resolution as well as in a variety of inflammatory conditions. Apart from its’ anti-secretory and anti-inflammatory characteristics, AF is an inhibitor of capillary leakage in intestine. It is not known whether the protein AF or the peptide AF-16 can ameliorate symptoms in sepsis. We hypothesized that AF-16 decreases the degree of hemodynamic instability, the need of fluid resuscitation, vasopressor dose and tissue edema in fecal peritonitis.

To test the hypothesis, we induced peritonitis and sepsis by injecting autologous fecal solution into abdominal cavity of anesthetized pigs, and randomized (in a blind manner) the animals to intervention (AF-16, n=8) or control (saline, n=8) group. After onset of hemodynamic instability (defined as mean arterial pressure < 60 mmHg maintained for > 5 minutes), resuscitation was initiated by an infusion of AF-16 or saline. We recorded respiratory and hemodynamic parameters hourly for twenty hours and collected post mortem tissue samples at the end of the experiment.

No differences between the groups were observed regarding hemodynamics, fluid balance, lung mechanics, gas exchange or histology. This experimental study suggests that AF-16 does not modulate sepsis symptoms in peritonitis induced sepsis.

## Introduction

Sepsis is defined as “life-threatening organ dysfunction caused by a dysregulated host response to infection” [1]. In septic shock profound circulatory and metabolic abnormalities contribute to an increase in mortality, with up to 40 % in-hospital mortality [1–3]. Requirement of vasopressor therapy to sustain a mean arterial pressure (MAP) > 65 mmHg in combination with persistent serum lactate level > 2 mmol/L after fluid resuscitation are the clinical hallmarks of septic shock [1]. Sepsis and septic shock are common and although sepsis mortality is decreasing [4], global estimates still count for more than 30 million cases of sepsis per year with 5.3 million potential fatalities [5].

Sepsis management is based on fluid resuscitation, broad-spectrum antibiotics, source control and vasoactive drugs [6]. Administration of intravenous fluid is fundamental to maintain adequate stroke volume and perfusion pressure, but as a consequence of fluid therapy patients often present with volume overload [7]. A positive fluid balance and volume overload is associated with longer stay in intensive care, prolonged use of mechanical ventilation and increased mortality [8–11].

Antisecretory factor (AF) is a 41 kDa protein detectable in most tissues [12]. The protein is secreted to plasma and becomes activated upon exposure to e.g. bacterial toxins [13]. AF has anti-secretory and anti-inflammatory properties [12,14,15]. The biologically active site of the protein is located in a 16-peptide fragment, AF-16, with the sequence VCHSKTRSNPENNVGL [16].

AF was first described as a potent inhibitor of intestinal hypersecretion in response to Cholera toxin [17]. It has since then been discovered to have multiple modulatory effects in altered fluid transport and edema formation/resolution [18–20] as well as in a variety of inflammatory conditions [14,21,22]. We have demonstrated in a previous study that AF-16 significantly reduced the fluid accumulation in the lungs in a porcine ventilator induced lung injury model [23]. AF is constitutively expressed in macrophages and is detectable in lymphoid organs, including gut-associated lymphoid tissue, spleen and thymus. The protein also appears to modulate proliferation of T cells [15]. Upon a pro-inflammatory stimulus AF expression is increased and the protein is redistributed from the perinuclear area to the cell surface [14,15]. This results in down-regulation of the immune response. AF is also an inhibitor of Cholera toxin induced capillary leakage [24].

Sepsis consists of a dysregulation of the fine-tuned balance between the pro- and anti-inflammatory systems. It is not known if AF or AF-16 could reverse shock symptoms in sepsis. We hypothesized that the peptide AF-16 could counteract circulatory instability in a porcine model of peritonitis induced sepsis, by reducing the inflammatory response (as disclosed by histopathology) and/or interstitial edema formation.

## Materials and methods

The study (protocol: http://dx.doi.org/10.17504/protocols.io.bdrsi56e) was approved by the Animal Ethics Committee in Uppsala (decision 5.8.18-01054/2017). The care of the animals strictly followed the National Institute of Health guide for the care and use of Laboratory animals (NIH publications No 8023, revised 1978) and all measures were taken to minimize suffering. The study was performed at the Hedenstierna Laboratory, Uppsala University, Sweden.

### Anesthesia and instrumentation

Sixteen pigs (*Sus scrofa domesticus*) (mean weight 27.3 +/− 2.4 kg) of mixed Swedish, Hampshire and Yorkshire breeds of both sexes, were sedated with Zoletil Forte (tiletamine and zolazepam) 6 mg/kg and Rompun (xylazine) 2.2 mg/kg i.m. A peripheral intravenous catheter was introduced in an ear vein. The animals were after 5-10 min placed supine and a bolus of fentanyl 5-10 μg/kg i.v. was administered, after which anesthesia was maintained with ketamine 30 mg/kg/h, midazolam 0.1-0.4 mg/kg/h and fentanyl 4 μg/kg/h, in glucose 2.5%. Esmeron (rocuronium) 2.5 mg/kg/h was added as muscle relaxant after adequate depth of anesthesia was assured by absence of reaction to painful stimulation between the front hooves. During the first hour thirty ml/kg/h of Ringer’s acetate was infused i.v. From the second hour until induction of peritonitis Ringer’s acetate was infused at a rate of 10 ml/kg/h.

After induction of anesthesia, the animals were tracheostomized, and a tube of eight mm internal diameter (Mallinckrodt Medical, Athlone, Ireland) was inserted in the trachea and connected to a ventilator (Servo I, Maquet, Solna, Sweden). Volume controlled ventilation was maintained with the following settings: tidal volume (V_T_) 8 ml/kg, respiratory rate (RR) 25/min, inspiratory/expiratory time (I:E) 1:2, inspired oxygen concentration (F_I_O_2_) 0.3 and positive end-expiratory pressure (PEEP) 8 cmH_2_O; V_T_, I:E and PEEP were maintained constant throughout the protocol. F_I_O_2_ was adjusted aiming at PaO_2_ >10 kPa. Respiratory rate was set at 25, but adjusted to keep PaCO_2_ <6,5 kPa.

A pulmonary artery catheter (Edwards Life-Science, Irvine CA, USA) for measurement of cardiac output (CO) and pulmonary artery pressures, and a triple lumen central venous catheter for fluid infusions were inserted via the right jugular vein. An arterial catheter for blood sampling and blood pressure measurement was inserted in the right carotid artery, and a PiCCO (pulse contour cardiac output) catheter (PV2015L20, Pulsion, Munich, Germany) was inserted in the right femoral artery for estimation of stroke volume variation (SVV) and extravascular lung water (EVLW). Blood gas analysis was executed on an ABL 3 analyzer, (Radiometer, Copenhagen, Denmark) and performed immediately after sampling. Hemoglobin and hemoglobin oxygen saturation was separately analyzed with a hemoximeter OSM 3 (Radiometer, Copenhagen, Denmark) calibrated for porcine hemoglobin.

A midline laparotomy was performed and the bladder catheterized for urinary drainage. Caecum was identified and a small incision made, feces was collected and the incision closed. A large-bore intra-peritoneal drain was inserted, and the abdominal incision closed.

### Study protocol

Preparation was followed by at least 30 min of stabilization, after which baseline measurements were performed (Fig 1). Fecal peritonitis was induced by peritoneal instillation of autologous feces (2 g/kg body weight in 200 ml warmed 5% glucose solution). The intraperitoneal drain was removed, and the abdominal wall closed. With the induction of fecal peritonitis the infusion of Ringer’s Acetate was discontinued.

**Fig 1.**
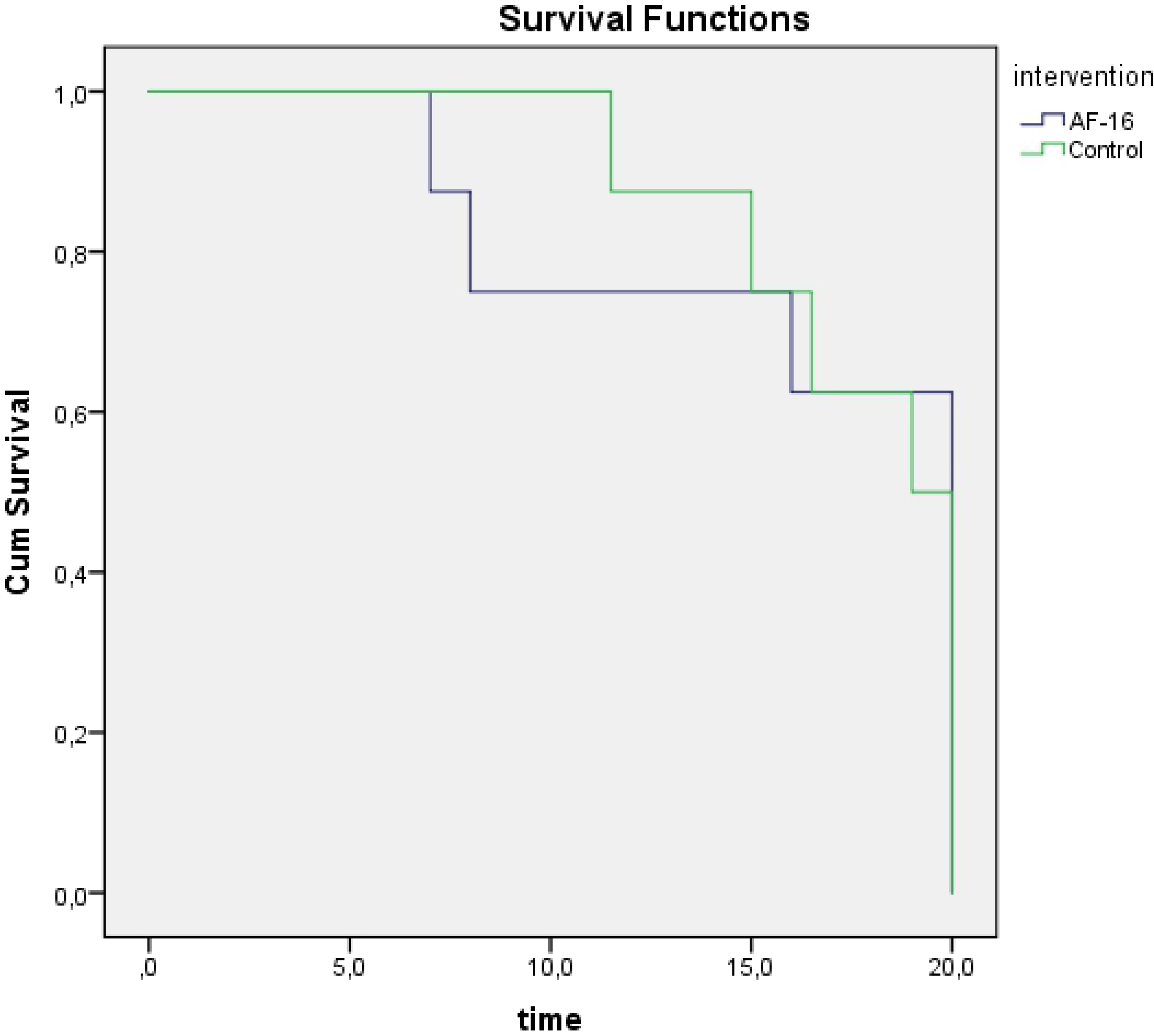
Experimental Time Line.

After peritonitis induction, animals were randomized to intervention with AF-16 (n=8) or control group (n=8), (block randomization: 4×4 sealed, opaque envelopes). The research team was blinded for the group allocation. Following the onset of hemodynamic instability (defined as MAP <60 mmHg for >5 min) the intervention group received an initial bolus of AF-16 (Batch No. 09431, KJ Ross Petersen ApS, Copenhagen, Denmark) 20 mg/kg (50 mg/ml in 0.9% saline), over duration of 10 minutes. The initial bolus dose was followed by an infusion of 40 mg/kg over 50 minutes. The control group received equal volumes of the vehicle (0.9% saline) instead. After four and eight hours the bolus dose was repeated (AF-16 or vehicle). Piperacillin/Tazobactam 2 gram every 8 hours i.v. and a protocolized resuscitation were initiated following established hemodynamic instability.

Both intervention and control groups were submitted to a protocolized resuscitation aiming at a MAP > 60 mmHg. Fluid resuscitation was initiated with Ringer’s Acetate 10 ml/kg/h. If signs of hypovolemia (SVV > 15% maintained for 10 min) a bolus of 150 ml Ringer’s Acetate was administered. Fluid boluses were repeated until SVV was stable < 15%. When SVV decreased to < 13% with MAP >60 mmHg, infusion was tapered down to 5 ml/kg/h, and if the animal was stable and SVV maintained < 13% the infusion was stopped. If signs of hypovolemia again appeared infusion was first started with 5 ml/kg/h then 10 ml/kg/h, then boluses of 150 ml were administered. In case of hypotension (MAP < 60 mmHg) without increased SVV, infusion of norepinephrine 5 ml/h (40 μg/ml) was started following a bolus of 1 ml (40 μg/ml), and increased stepwise by 5 ml/h. Glucose 30 % infusion was administered, aiming at blood glucose 5-10 mmol/L, starting with 0.5 ml/kg/h. If b-glucose > 10 mmol/L an insulin infusion 1E/ml was started with 1 ml/h.

We performed blood gas analyses at baseline, after onset of shock and every hour for the following twenty hours duration of the experiment. At the same time points, hemodynamic parameters (systemic and pulmonary pressures, CO, heart rate), respiratory parameters (F_I_O_2_, SaO_2_, ETCO_2_, plateau pressure, dynamic and static compliance) and hourly urine output were measured. Every three hours EVLW was measured and mixed venous blood gas analysis performed. Stroke volume variation (SVV) was monitored continuously in order to guide fluid administration.

The animals were euthanized with 100 mmol KCl i.v. at the end of the experiment under deep anesthesia. Thereafter the chest wall was opened. Lung tissue samples were collected from both lungs from the following regions: apical-medial, medial-medial, caudal-dorsal, caudal-medial and caudal-ventral. Samples were also taken from heart, liver, kidney, intestine and skin. The samples were immediately immersed in 10% buffered formalin. A veterinary pathologist who was blinded for the group allocation evaluated the samples histologically. Wet-to-dry ratio was measured in the above mentioned lung regions from the right lung. Samples were weighed, and dried in an oven, at 50° C, until the weight did not differ between two measurements.

### Statistical analysis

The Mead Resource Equation was used to determine sample size [25]. We used the Shapiro-Wilk test to test the data for normality. We compared groups with the two-tailed Student’s t-test, Mann-Whitney U-test, or the Kruskal-Wallis test. Two-way repeated measures ANOVA was used to compare differences within and between the groups over time. Tukey post hoc test was applied when appropriate. Last observation carried forward was used as imputation of missing data because of early deaths. The data are expressed as mean +/− SD or median (interquartile range) when appropriate. The statistical analyses were conducted by SPSS v. 20.0.0 software (SPSS, Inc., Chicago, IL, USA). A *p*-value of < 0.05 was considered to be statistically significant.

## Results

The two groups were comparable at baseline regarding hemodynamics and respiratory parameters (Table 1). Mean time from peritonitis induction to onset of hemodynamic instability was 4.5 +/− 2.2 and 4.9 +/− 1.2 hours in treatment and control groups, respectively.

**Table 1.** Measurements at baseline.

|  | **AF-16** | **Control** |
| --- | --- | --- |
| **MAP (mmHg)** | 74 +/- 13 | 71 +/- 12 |
| **HR (BPM)** | 82 +/- 11 | 90 +/- 10 |
| **CO (l/min)** | 2.4 +/- 0.7 | 2.7 +/- 0.7 |
| **MPAP (mmHg)** | 19 +/- 3 | 18 +/- 3 |
| **EVLW (ml)** | 360 +/- 90 | 290 +/- 40 |
| **SVV (%)** | 10 +/- 4 | 8 +/- 4 |
| **PaO2/FiO2 (kPa)** | 60 +/- 3 | 61 +/- 5 |
| **Static compliance (ml/cmH<sub>2</sub>O)** | 34 +/- 6 | 32 +/- 5 |

Values expressed as mean +/- SD. MAP (mean arterial pressure), HR (heart rate), CO (cardiac output), MPAP (mean pulmonary arterial pressure), EVLW (extra vascular lung water), SVV (stroke volume variation) and PaO2/FiO2 (the arterial oxygen tension/ inspired oxygen tension).

Nine out of the sixteen animals survived the experiment until euthanasia (20 hours), while four and three animals died during the 20-hours observation period in treatment and control groups, respectively (Fig 2). There was no statistically significant difference in survival between intervention and control groups.

**Fig 2.**
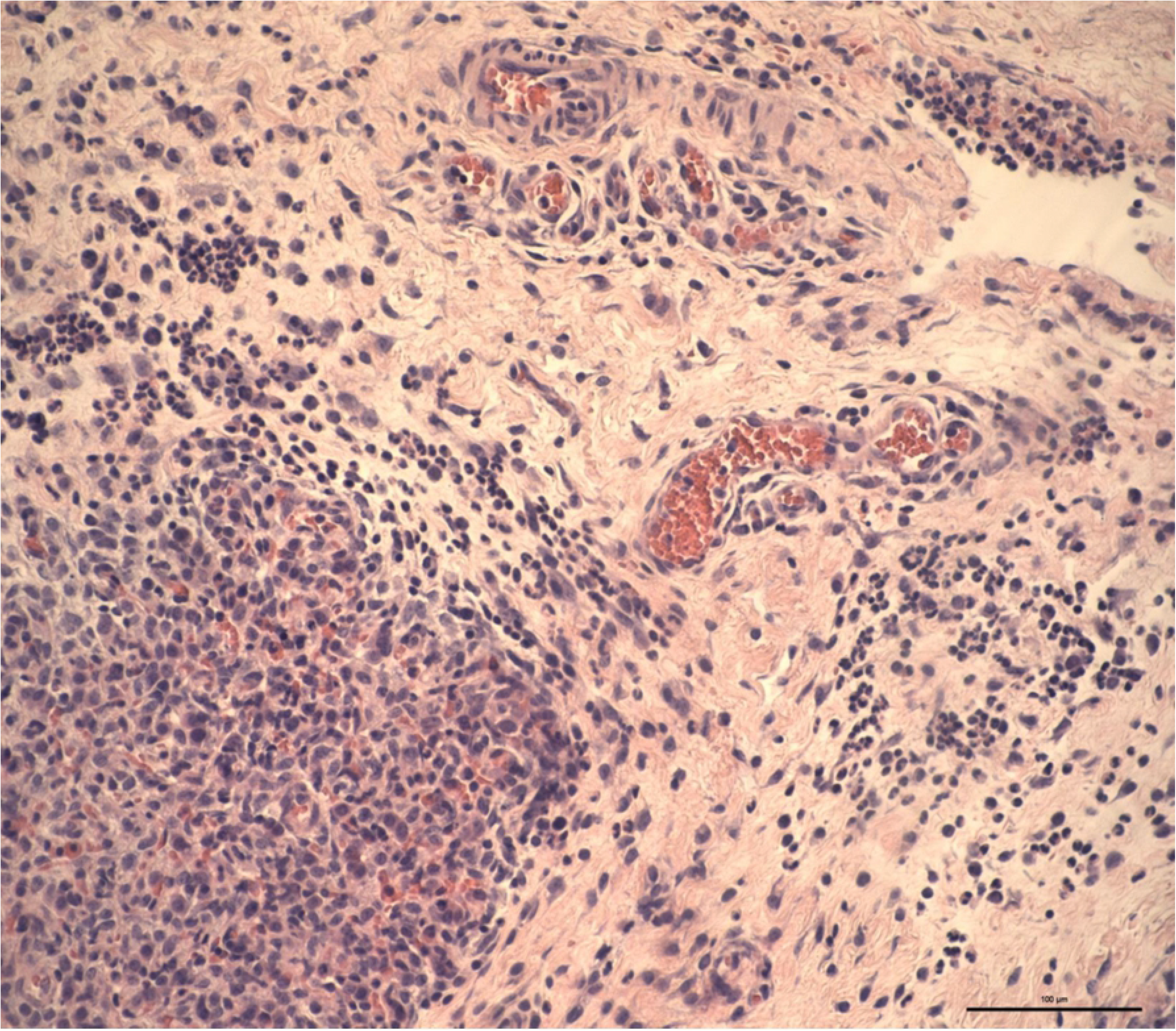
Kaplan-Meyer analysis of survival.

### Gas exchange and lung mechanics

#### Gas exchange

After established hemodynamic instability both groups presented with a decline in oxygenation (PaO_2_/F_I_O_2_ ratio). The AF-16 group went from 60 +/− 3.0 kPa at baseline, to 33 +/− 13.9 kPa at the end of the protocol, the control group went from 61 +/− 4.7 kPa, to 27 +/− 15.7 kPa at the same time points (Table 2). There was no statistically significant difference in oxygenation between the intervention and control groups (two way ANOVA F (2, 54) = 0.093, *p* = 1) as a function of time. Respiratory rate was adjusted to keep PaCO_2_ under 6.5 kPa (Table 2).

**Table 2.** Respiratory parameters.

|  | Group | Baseline | S0 | End |
| --- | --- | --- | --- | --- |
| **PaO<sub>2</sub>/F<sub>I</sub>O<sub>2</sub> (kPa)** | AF-16 | 60 +/- 3.0 | 49 +/- 3.5 | 33 +/-13.9 |
|  | Control | 61 +/- 4.7 | 49 +/- 6.9 | 27 +/- 15.7 |
| **Driving Pressure (mmHg)** | AF-16 | 7 +/- 1.2 | 9 +/- 1.5 | 18 +/- 2.6 |
|  | Control | 8 +/- 3.2 | 9 +/- 2.8 | 20 +/- 10.8 |
| **Static compliance (ml/cmH<sub>2</sub>O)** | AF-16 | 34 +/- 6.3 | 26 +/-2.4 | 15 +/- 2.9 |
|  | Control | 32 +/- 5.4 | 27 +/-5.7 | 15 +/- 5.1 |
| **Dynamic compliance (ml/cmH<sub>2</sub>O)** | AF-16 | 28 +/- 4.2 | 23 +/- 2.9 | 12 +/- 3.0 |
|  | Control | 26 +/- 6.1 | 22 +/- 4.7 | 11 +/- 4.5 |
| **Saturation (%)** | AF-16 | 96 +/- 0.3 | 94 +/- 1.4 | 91 +/- 5.9 |
|  | Control | 97 +/- 0.9 | 94 +/- 1.4 | 85 +/- 10.9 |
| **MPAP (mmHg)** | AF-16 | 19 +/- 2.8 | 22 +/- 3.0 | 29 +/- 5.9 |
|  | Control | 18 +/- 2.7 | 23 +/- 3.6 | 28 +/- 5.6 |
| **PaCO<sub>2</sub> (kPa)** | AF-16 | 5.5 +/- 0.4 | 5.9 +/- 0.1 | 4.9 +/- 1.5 |
|  | Control | 5.2 +/- 0.5 | 5.8 +/- 0.4 | 6.5 +/- 1.2 |

Values expressed as means +/- SD. PaO<sub>2</sub>/F<sub>I</sub>O<sub>2</sub> (the arterial oxygen tension/ inspired oxygen tension), MPAP (mean pulmonary arterial pressure), PaCO<sub>2</sub> (arterial CO<sub>2</sub> tension).

#### Lung mechanics

Static compliance decreased from 34 +/− 6.3 ml/cm H_2_O at baseline in the intervention group, to 15 +/− 2.9 ml/cm H_2_O at the end of the protocol and from 32 +/− 5.4 ml/cm H_2_O to 15 +/− 5.1 ml/cm H_2_O in the control group (Table 2), (two-way ANOVA, F (21,252) = 0.145, *p* = 1.00). Dynamic compliance and driving pressure changed comparably in both groups during the length of the experiment (Table 2).

### Hemodynamic parameters

#### Extravascular lung water (EVLW) and Stroke Volume Variation (SVV)

There was no statistically significant difference in EVLW evolution between intervention and control groups as a function of time (two-way ANOVA, F (7,87) = 0.77, *p* = 0.614). EVLW increased from 360 +/− 90 ml at baseline to 550 +/− 370 ml at the end of the observation period, and from 290 +/− 40 ml to 450 +/− 300 ml in the intervention and control groups, respectively (Table 3). Neither was there any statistically significant difference between groups as a function of time regarding SVV (Table 3).

**Table 3.** Hemodynamic parameters.

|  | Group | Baseline | S0 | End |
| --- | --- | --- | --- | --- |
| MAP (mmHg) | AF-16 | 74 +/- 12.5 | 57 +/- 2.9 | 59 +/- 17.6 |
|  | Control | 71 +/- 11.6 | 57 +/- 2.7 | 58 +/- 14.5 |
| HR (BPM) | AF-16 | 82 +/- 11 | 159 +/- 21 | 117 +/- 34 |
|  | Control | 90 +/- 10 | 139 +/- 32 | 112 +/- 23 |
| CO (l/min) | AF-16 | 2.4 +/- 0.7 | 2.5 +/- 0.2 | 2.9 +/- 2.0 |
|  | Control | 2.7 +/- 0.7 | 2.1 +/- 0.3 | 2.9 +/- 1.2 |
| pH | AF-16 | 7.44 +/- 0.03 | 7.36 +/- 0.03 | 7.33 +/- 0.18 |
|  | Control | 7.47 +/- 0.04 | 7.39 +/- 0.04 | 7.24 +/- 0.20 |
| Lactate (mmol/l) | AF-16 | 3.0 +/- 1.2 | 2.6 +/- 1.0 | 3.6 +/- 4.0 |
|  | Control | 2.6 +/- 1.0 | 2.2 +/- 0.8 | 3.9 +/- 4.0 |
| Hb (g/l) | AF-16 | 87 +/- 7.7 | 119 +/- 12 | 94 +/- 10 |
|  | Control | 85 +/- 5.3 | 122 +/- 5 | 100 +/- 6 |
| SVV (%) | AF-16 | 10 +/- 3.6 | 21 +/- 7.7 | 15 +/- 8.1 |
|  | Control | 8 +/- 4.0 | 14 +/- 2.0 | 16 +/- 6.4 |
| EVLW (ml) | AF-16 | 360 +/- 90 | 400 +/- 14 | 550 +/- 370 |
|  | Control | 290 +/- 40 | 280 +/- 40 | 450 +/- 300 |
| ERO<sub>2</sub> (kPa) | AF-16 | 58.6 +/- 12.5 | 55.4 +/- 6.6 | 49.1 +/- 12.8 |
|  | Control | 49.4 +/- 9.3 | 51.7 +/- 9.0 | 44.3 +/- 8.6 |

Values as mean +/- SD. MAP (mean arterial pressure), HR (heart rate), CO (cardiac output), Hb (Hemoglobin concentration), SVV (stroke volume variation), EVLW (extra vascular lung water), and ERO<sub>2</sub> (the oxygen extraction ratio).

#### Mean arterial blood pressure and heart rate

The onset of hemodynamic instability was defined as mean arterial pressure under 60 mmHg, both intervention and control groups presented with increases in heart rate at this stage of the experiment (Table 3). There was no statistically significant difference between groups regarding heart rate throughout the observation period (Two way ANOVA, F(21,252) = 0.765, *p* =0.761). Onset of hemodynamic instability was also accompanied by an increase in hemoglobin concentration in both groups, while no statistically significant difference between groups was detected (Two way ANOVA, F(21,253) = 0.214, *p* = 1.00). The two groups did not differ in a statistically significant way in either lactate, pH or oxygen extraction ratio (Table 3).

#### Fluid balance

There was no statistically significant difference between groups in fluid requirements, urinary output, fluid balance (these parameters described as ml/kg/h of sepsis duration), norepinephrine consumption (μg/kg/h of sepsis duration) nor percentage body weight gain (kg body weight before and after experiment) (Table 4).

**Table 4.** Fluid balance.

|  | **AF-16** | **Control** |
| --- | --- | --- |
| **Fluid requirement**<br>(ml/kg/h sepsis) | 17 +/-10 | 15 +/- 4 |
| **Urinary output**<br>(ml/kg/h sepsis) | 1,3 +/- 1 | 1,5 +/- 1 |
| **Fluid balance**<br>(ml/kg/h sepsis) | +16 +/- 10 | +14 +/- 4 |
| **Norepinephrine**<br>($\mu\text{g/kg/h}$ sepsis) | 37.5 +/- 32.6 | 32.3 +/- 24.6 |
| **% weight gain** | 33 +/- 22 | 51 +/- 26 |

#### Wet-to-dry ratio

Wet-to-dry ratio at the end of the experiment did not differ in a statistically significant manner between intervention and control groups. Samples from lung, skin, intestine, heart (left ventricle), kidney and liver were analyzed. Lung samples were analyzed separately and pooled together. Skin had the lowest water content, kidney the highest (Table 5).

**Table 5.** Wet-to-dry ratio.

|  | **AF-16** | **Control** |
| --- | --- | --- |
| **Lung w/d** | 1,06 +/- 0,01 | 1,06 +/- 0,01 |
| **Skin w/d** | 1,02 +/- 0,01 | 1,01 +/- 0,01 |
| **Intestine w/d** | 1,08 +/- 0,04 | 1,08 +/- 0,04 |
| **Heart w/d** | 1,08 +/- 0,02 | 1,07 +/- 0,03 |
| **Kidney w/d** | 1,13 +/- 0,05 | 1,10 +/- 0,03 |
| **Liver w/d** | 1,08 +/- 0,02 | 1,08 +/- 0,02 |

Values expressed as mean +/- SD.

### Histology

Abnormal lesions were found most commonly in the lungs and the intestine. The intensity of lesions was graded in a semi quantitative way, based on the numbers of inflammatory cells and the extension and distribution of the cell infiltrates/lesions. Inflammatory cell exudates in lung samples included neutrophils, monocytes and macrophages. Leucocytes were increased in the interstitium, vessels and perivascular space (Fig 3a and 3b).

**Figs 3a and 3b.**
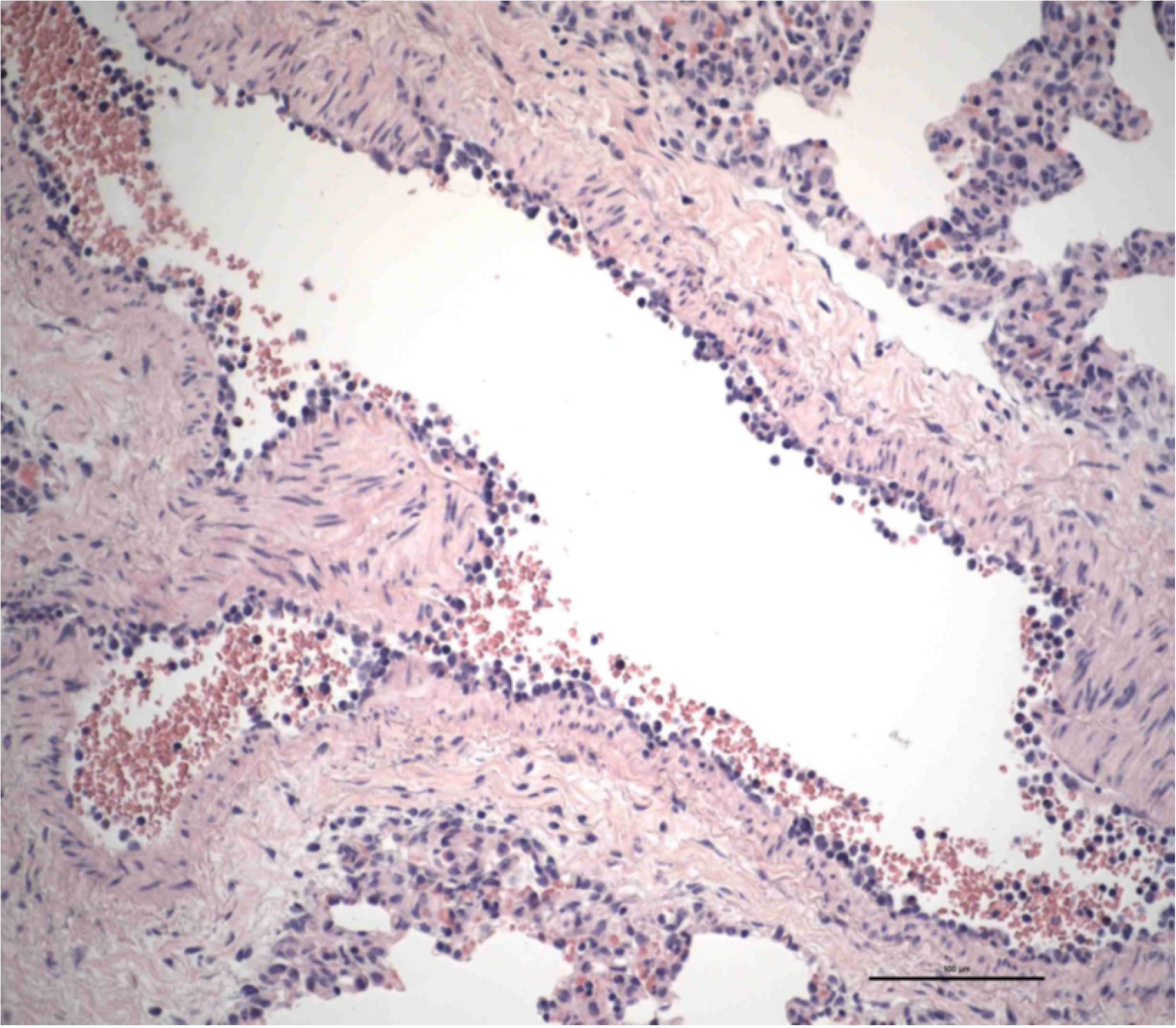
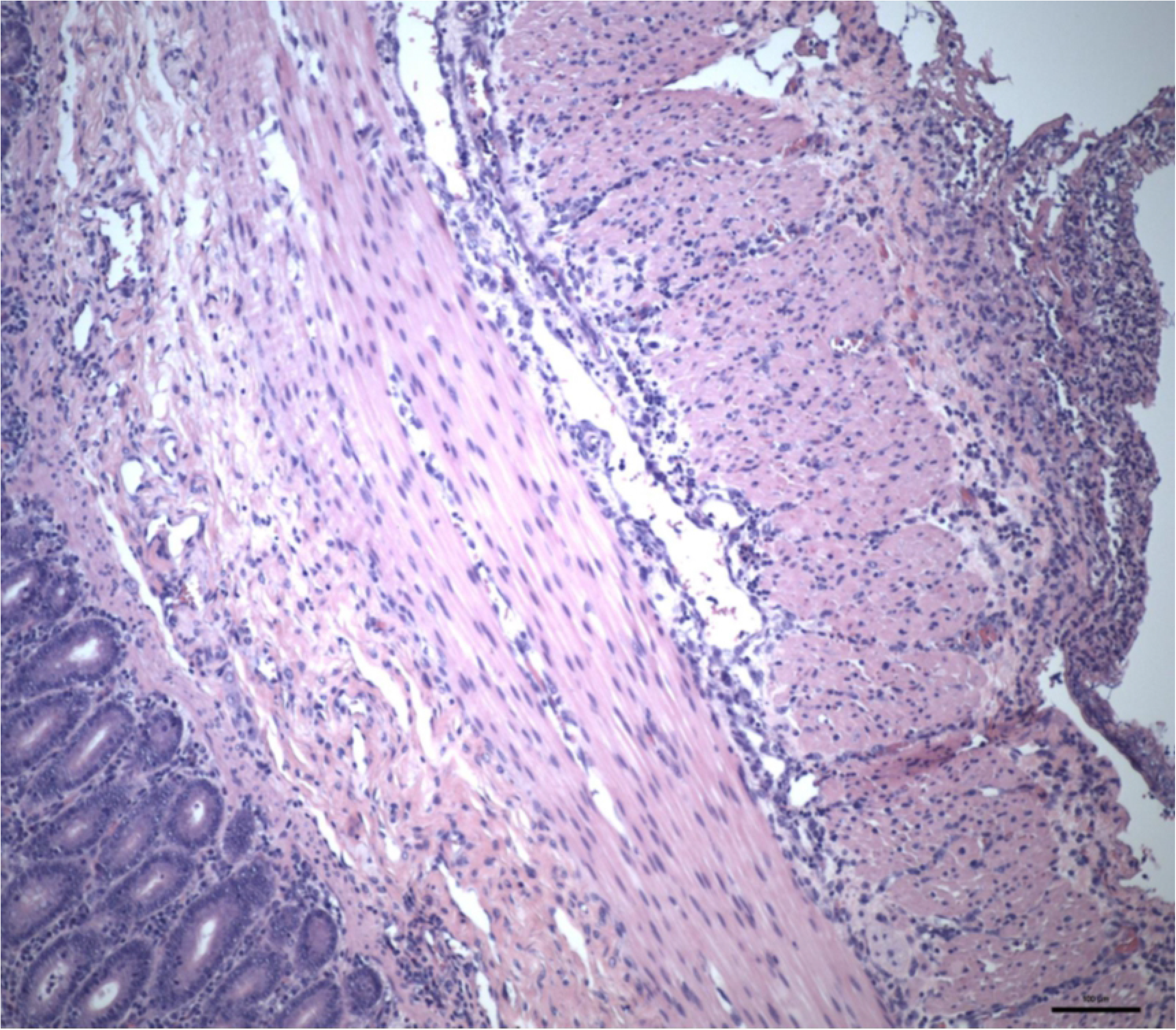
Histology of lung samples, AF-16 vs control. **3a** Intense inflammatory cell reaction shows leukocytes rich in polymorphs in the alveoli (down, at left) and in the capillaries in the interlobular septum (AF-16). **3b** Bronchial vessel shows leukocytes adhering at the endothelium. It could be an early stage in the process of leukocyte migration through the vessel wall, but leukocytes seem to remain in the intima which is suggestive of endoarteriolitis, which could be predisposing for thrombosis (control).

**Figs 4a and 4b.**
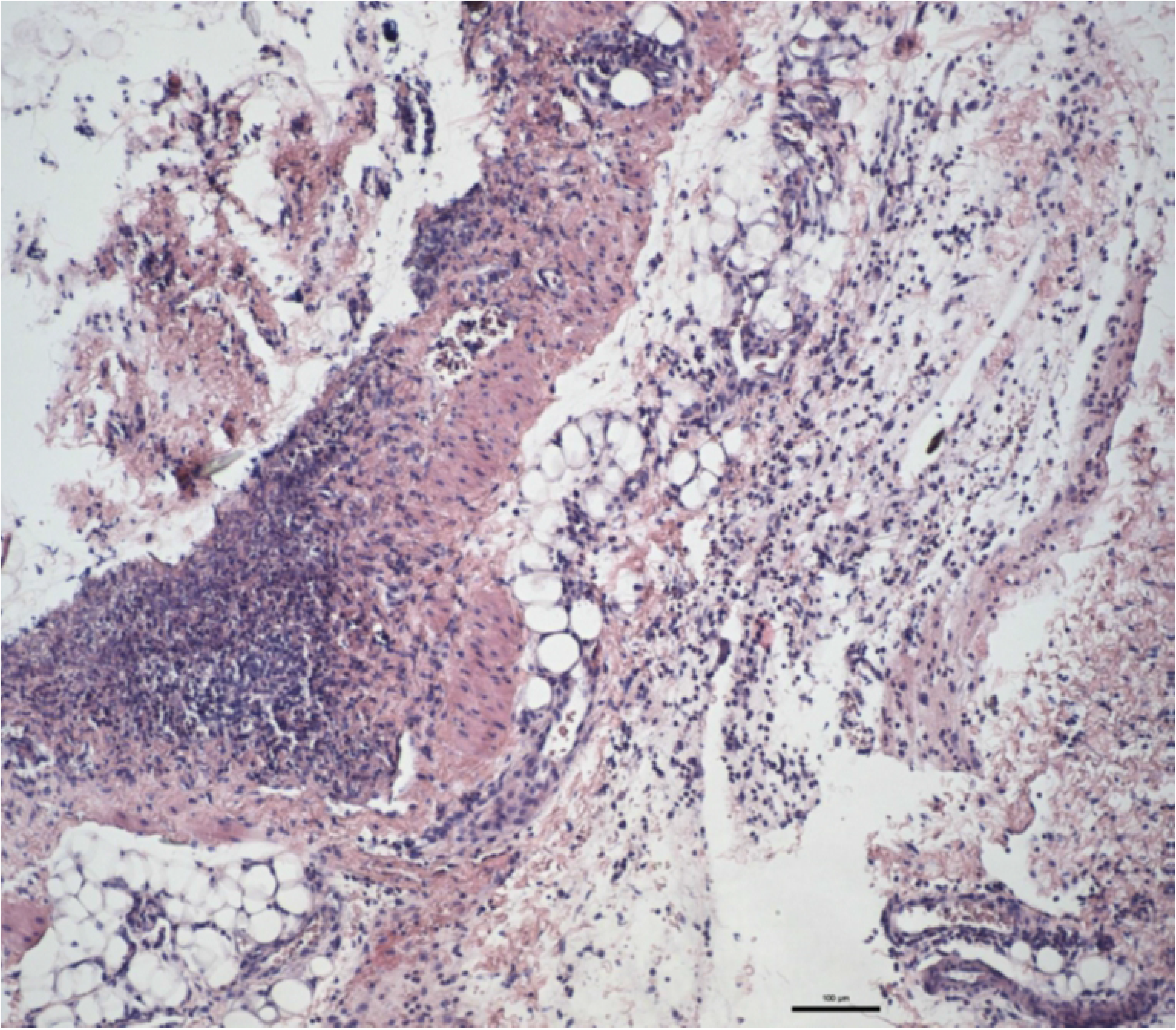
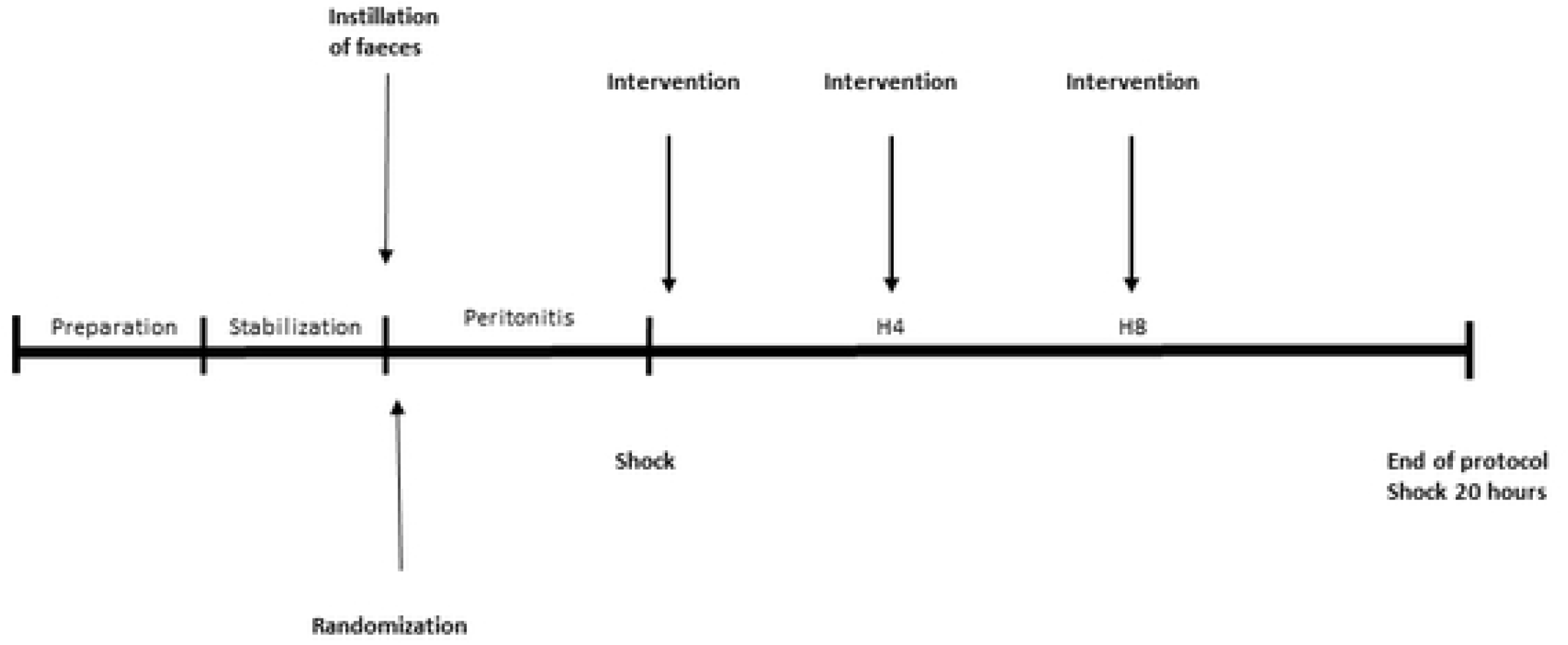
Histology of intestine and mesenterium. 4a the mucosa is down at left and the serosa is at right. The serosa shows a rich fibrinopurulent exudate, consistent with peritonitis. Leukocytes between the smooth muscle layers also are visible (AF-16). 4b Mesenterium. Necrosis and intense inflammatory reaction in the fat and connective tissues, consistent with peritonitis (control).

Vessels often displayed prominent endothelial cells and leucocytes were found in the process of margination and migration through the vessel wall. Many lung samples showed edema, hemorrhages and recently originated micro-thrombi in small-sized vessels commonly blocking the lumen; sometimes with adjacent alveolar areas with congested septal capillaries, hemorrhages and pyknotic cells suggestive of necrosis. There was no statistically significant difference between intervention and control groups regarding inflammation or edema in lung samples (Table 6).

**Table 6.** Lung Histology.

|  | **Leukocytes,**<br>**AF-16** | **Leukocytes,**<br>**Control** | **Atelectasis,**<br>**AF-16** | **Atelectasis,**<br>**Control** | **Edema, AF-**<br>**16** | **Edema,**<br>**Control** |
| --- | --- | --- | --- | --- | --- | --- |
| **AMR** | 1.5 (0-3) | 2.5 (1-3) | 0.5 (0-3) | 0.5 (0-4) | 2 (0-4) | 3 (0-4) |
| **MMR** | 1.5 (0-4) | 1.5 (1-4) | 0.5 (0-4) | 1 (0-3) | 1.5 (0-4) | 2 (0-4) |
| **CMR** | 1.5 (0-3) | 3 (1-4) | 4 (2-4) | 3.5 (2-4) | 2.5 (0-4) | 2.5 (1-4) |
| **CDR** | 1.5 (0-2) | 3 (1-4) | 4 (2-4) | 4 (2-4) | 2.5 (0-2) | 3 (1-4) |
| **CVR** | 1.5 (0-2) | 2.5 (0-4) | 4 (3-4) | 4 (3-4) | 2.5 (1-4) | 3.5 (1-4) |

Values expressed as median (min-max). No statistically significant difference between groups, statistics presented for pooled data in the following order: leucocytes, atelectasis and edema (Kruskal-Wallis $p = 0.169$ , Kruskal-Wallis $p = 0.672$ , Kruskal-Wallis $p = 0.751$ ).

The intestines showed severe acute degenerative and necrotic changes in the mucosa. Some samples of intestine showed signs of transmural inflammation. There was no difference between intervention and control groups regarding signs of inflammation in samples of intestine, (Median 3, min 0, max 4 in both groups) (Kruskal-Wallis, *p* = 0.321). (Figs 5 and 6).

There were few signs of lesions (vacuoles, inflammation) in samples of heart, liver and kidney. No lesions were detected in skin biopsies.

## Discussion

In this experimental study of peritonitis induced sepsis, intervention with the anti-secretory and anti-inflammatory peptide AF-16 did not yield any reversal of sepsis symptoms as reflected in signs of inflammation, fluid balance, hemodynamics, tissue edema, norepinephrine consumption, gas exchange or respiratory mechanics.

We used a model of fecal peritonitis induced sepsis, previously described by Correa et al. [26]. Prior t the main protocol we performed a pilot study of four animals, in which all 4 animals died before finishing the protocol of fulminant shock symptoms (S7 Table, supplemental data). The animals of the pilot study presented with sepsis after a mean duration of peritonitis of 4.25 +/− 0.5 hours, mean survival 11.5 +/− 4.0 hours. Although the onset of hemodynamic instability (MAP <60 mmHg for more than five minutes) was the same in the sixteen animals included in the main series, sepsis was not as homogenously severe, and nine animals survived until euthanasia.

This model of sepsis has features in common with peritonitis induced sepsis in patients. The animals received autologous feces in the peritoneum to mimic intestinal perforation. Despite the prompt identification of hemodynamic instability in the absence of intravenous fluid administration, followed by immediate administration of antibiotics and fluid resuscitation, the mortality was substantial, 44% of the animals died before finishing the observation period of twenty hours.

The definition of septic shock in humans include an increased serum lactate > 2 mmol/L despite adequate fluid resuscitation. In this study we did not observe any statistically significant hyperlactatemia. Oxygen extraction ratio however declined significantly in both groups as a sign of either diminished oxygen demand or inefficient utilization of oxygen in the tissues.

Peritonitis induced impairment of gas exchange and lung mechanics in the current study were similar to Acute Respiratory Distress Syndrome (ARDS) in humans. At the end of the protocol all but three animals fulfilled oxygenation criteria for ARDS. This decrease in oxygenation was accompanied with a significant decline in both static and dynamic compliance and an increase in driving pressure with predefined tidal volumes.

There was no statistically significant difference in EVLW evolution between the groups during the experiment, all except three animals (one from intervention group and two control animals) did manifest an increase in EVLW, ranging from 7 % to 279 %.

In a previous study [23] we examined the potential effect of AF-16 on resolution of pulmonary edema in a model of ventilator induced lung injury, consisting of lung lavages and injurious ventilation. In that study a statistically significant reduction of EVLW in the intervention group was found, as an isolated finding. That finding was not reproduced in the present study, and although all animals did not respond with an increase in EVLW, leaving out the “EVLW non responders” in post-hoc analysis did not yield a different outcome.

The endogenous protein AF, and its active sequence AF-16, counteract edema and abnormal fluid flux [17–20,27,28]. In addition, AF protein/peptide exerts anti-inflammatory properties in a variety of conditions [15,21,22]. Neither AF nor AF-16 affect healthy tissue [28].

In models of edema and increased interstitial fluid pressure AF-16 has an early effect [18,28]. In a study by Jennische et al, intranasal administration of AF-16 reduced ICP after 15 min, but no effect on inflammatory response in brain could be discerned [19]. The endogenous AF response to an inflammatory stimulus is considered to be slower. Exposure to pro-inflammatory stimulus in form of LPS or IFN-γ results in an increase in AF expression and redistribution from perinuclear area to cell surface over a time period of several days, expression peaks with severity of disease and thereafter returns to baseline. It has previously been speculated that AF plays its main role in modifying the immune reponse in the resolution phase of an inflammatory reaction, rather than at the beginning of an immunological response [12,14,15]. AF-activity is low in health and in chronic inflammatory conditions, and therefor chronic inflammation might benefit better from treatment with AF/AF-16 than acute conditions [13].

In this study AF-16 was given in repeated doses, the initial dose being three times higher than in our previous ventilator induced lung injury (VILI) model [23]. We cannot rule out that the intervention with AF-16 would be more effective at an even higher or continuous dose, as AF in plasma has a rapid turnover rate [12], or that an effect could have been observed in a less severe shock state. Moreover, the number of animals studied was limited, so minor changes between the groups might not have been noticed. We cannot rule out the possibility that AF or AF-16 could be effective in a later stage of sepsis.

This study has limitations. No animal model reproduces the full picture of sepsis/septic shock in humans. The biological heterogeneity in sepsis patients, with differences in age, comorbidities, medications and different sources of infection adds to the complexity of the syndrome. This complexity cannot be fully represented in an animal model. In the present peritonitis/sepsis model the pigs are healthy prior to the experiment. One must also accept the possibility of interspecies variability in intestinal flora and host response to both infection and intervention.

We conclude that, contrary to our hypothesis, in this pilot study in a porcine experimental model of fecal peritonitis and sepsis we could not detect any differences between intervention and control groups regarding reversal of shock symptoms, gas exchange or respiratory mechanics. The results suggest that AF-16 does not ameliorate sepsis symptoms.

## References

1. Singer M, Deutschman CS, Seymour CW, Shankar-Hari M, Annane D, Bauer M., et al. The Third International Consensus Definitions for Sepsis and Septic Shock (Sepsis-3). JAMA. 2016;315:801–810.

2. Seymour CW, Liu VX, Iwashyna TJ, Brunkhorst FM, Rea. TD, Scherag A, et al. Assessment of Clinical Criteria for Sepsis For the Third International Consensus Definitions for Sepsis and Septic Shock (Sepsis-3). JAMA. 2016;315:762–774.

3. Shankar-Hari M, Phillips GS, Levy ML, Seymour CW, Liu VX, Deutschman CS, et al. Developing a New Definition and Assessing New Clinical Criteria for Septic Shock For the Third International Consensus Definitions for Sepsis and Septic Shock (Sepsis-3). JAMA. 2016;315:775–787.

4. Prescott H, Angus DC. Enhancing Recovery From Sepsis A Review. JAMA. 2019;319:62–75.

5. Fleischmann C, Scherag A, Adhikari NKJ, Hartog CS, Tsaganos T, Schlattmann P, et al. Assessment of Global Incidence and Mortality of Hospital-treated Sepsis. Am J Respir Crit Care Med. 2016;193:259–272.

6. Rhodes A, Evans LE, Alhazzani W, Levy MM, Antonelli M, Ferrer R, et al. Surviving Sepsis Campaign : International Guidelines for Management of Sepsis and Septic Shock : 2016. Intensive Care Med. 2017;43:304–377.

7. Mitchell KH, Carlbom D, Caldwell E, Leary PJ, Himmelfarb J, Hough CL. Volume Overload : Prevalence, Risk Factors, and Functional Outcome in Survivors of Septic Shock. Ann Am Thorac Soc 2015;12:1837–1844.

8. Acheampong A, Vincent J. A positive fluid balance is an independent prognostic factor in patients with sepsis. Crit. Care 2015;19: 1–7.

9. Malbrain MLNG, Marik PE, Witters I, Cordemans C, Kirkpatrick AW, Roberts DJ, et al. Fluid overload, de-resuscitation, and outcomes in critically ill or injured patients: a systematic review with suggestions for clinical practice. Anaesthesiol. Intensive Ther. 2014;46:361–380.

10. Oliveira FSV, Freitas FGR, Ferreira EM, Castro I, Bafi AT, Azevedo ACP, et al. Positive fluid balance as a prognostic factor for mortality and acute kidney injury in severe sepsis and septic shock. J. Crit. Care. 2015;30:97–101.

11. You JW, Lee SJ, Kim YE, Cho YJ, Jeong YY, Kim HC, et al. Association between weight change and clinical outcomes in critically ill patients. J. Crit. Care. 2013;28:923–927.

12. Lange S, Lönnroth I. The Antisecretory Factor: Synthesis, Anatomical and Cellular Distribution, and Biological Action in Experimental and Clinical Studies. Int. Rev. Cytol. 2001;210: 39–74.

13. Johansson E, Lönnroth I, Jonson I, Lange S, Jennische E. Development of monoclonal antibodies for detection of Antisecretory Factor activity in human plasma. J. Immunol. Methods 2009;342:64–70.

14. Davidson TS, Hickey WF. Antisecretory factor expression is regulated by inflammatory mediators and influences the severity of experimental autoimmune encephalomyelitis. J. Leucoc. Biol. 2004;74:835–844.

15. Davidson TS, Hickey WF. Distribution and immunoregulatory properties of antisecretory factor. Lab. Investig. 2004;84:307–319.

16. Johansson E, Lange S, Lönnroth I. Identification of an active site in the antisecretory factor protein. Biochim. Biophys. Acta 1997;1362:177–182.

17. Lönnroth I, Lange S. Purification and characterization of the antisecretory factor: a protein in the central nervous system and in the gut which inhibits intestinal hypersecretion induced by cholera toxin. Biochim. Biophys. Acta 1986;883:138–144.

18. Clausen F, Hansson HA, Raud J, Marklund N. Intranasal Administration of the Antisecretory Peptide AF-16 Reduces Edema and Improves Cognitive Function Following Diffuse Traumatic Brain Injury in the Rat. Front Neuro 2017;8:1–14.

19. Jennische E, Bergström T, Johansson M, Nyström K, Tarkowski A, Hansson HA. et al. The peptide AF-16 abolishes sickness and death at experimental encephalitis by reducing increase of intracranial pressure. Brain Res. 2008;1227:189–197.

20. Johansson E, Al-Olama M, Hansson HA, Lange S, Jennische E. Diet-induced antisecretory factor prevents intracranial hypertension in a dosage-dependent manner. Br. J. Nutr. 2013; 109:247–2252.

21. Björck S, Bosaeus I, Ek E, Jennische E, Lönnroth I, Johansson E, et al. Food induced stimulation of the antisecretory factor can improve symptoms in human inflammatory bowel disease: A study of a concept. Gut 2000;46:824–829.

22. Eriksson A, Shafazand M, Jennische E, Lange S. Effect of antisecretory factor in ulcerative colitis on histological and laborative outcome: a short period clinical trial. Scand. J. Gastroenterol. 2003;5521:1045–1049.

23. Tenhunen AB, Massaro F, Hansson HA, Feinstein R, Larsson A, Larsson A, et al. Does the Antisecretory Factor reduce lung edema in experimental ARDS. Ups J Med Sci. 2019;124:246–253

24. Lange S, Delbro DS, Jennische E, Johansson E, Lönnroth I. Recombinant or Plasma-Derived Antisecretory Factor Inhibits Cholera Toxin-Induced Increase in Evans Blue Permeation of Rat Intestinal Capillarie s. Dig. Dis. Sci. 1998;43:2061–2070.

25. Mead R. The Non-Orthogonal Design of Experiments. J R Stat Soc. 1990;153(2):151–201.

26. Corrêa TD, Vuda M, Takala J, Djafarzadeh S, Silva E, Jakob SM. Increasing mean arterial blood pressure in sepsis : effects on fluid balance, vasopressor load and renal function. Crit. Care 2013;17:R21.

27. Hansson HA, Al-Olama M, Jennische E, Gatzinsky K, Lange S. The Peptide AF-16 and the AF Protein Counteract Intracranial Hypertension. Acta Neurochir. Suppl. 2012;114:377–382.

28. Al-Olama M, Wallgren A, Andersson B, Gatzinsky K, Hultborn R, Karlsson-Parra A, et al. The peptide AF-16 decreases high interstitial fluid pressure in solid tumors. Acta Oncol. 2011;50:1098–1104.

